# Dissociable effects of urgency and evidence accumulation during reaching revealed by dynamic multisensory integration

**DOI:** 10.1101/2023.12.15.571806

**Authors:** Anne H. Hoffmann, Frédéric Crevecoeur

## Abstract

When making perceptual decisions, humans combine information across sensory modalities dependent on their respective uncertainties. However, it remains unknown how the brain handles multisensory integration during movement, and which factors besides sensory uncertainty might influence the contribution of different modalities. We performed two reaching experiments on healthy adults to investigate whether movement corrections to combined visual and mechanical perturbations scale with visual uncertainty. To describe the dynamics of multimodal feedback responses, we further varied movement speed and duration of visual feedback during the movement. The results of our first experiment (N=16, 11 females) show that the contribution of visual feedback decreased with uncertainty. Interestingly, we observed a transient phase during which visual feedback responses were stronger during faster movements. In a follow-up experiment (N=16, 10 females), we found that the contribution of vision increased more quickly during slow movements when we presented the visual feedback for a longer time. Using an optimal feedback control model, we show that the increased response to visual feedback during fast movements can be explained by an urgency-dependent increase in control gains. Further, the fact that viewing duration increased the visual contributions suggests that the brain indeed performs a continuous state-estimation as expected in the optimal control model featuring a Kalman filter. Hence, both uncertainty and urgency determine how the sensorimotor system responds to multimodal perturbation during reaching control. We highlight similarities between reaching control and decision-making, both of which appear to be influenced by the accumulation of sensory evidence as well as response urgency.

**Significance statement:** The exact time course of multisensory integration during movement, along with the factors that influence this process, still requires further investigation. Here, we tested how visual uncertainty, movement speed, and visual feedback duration influence corrective movements during reaching with combined visual and mechanical perturbations. Using an optimal feedback control model, we illustrate that the time course of multimodal feedback responses follows the predictions of a Kalman filter which continuously weighs sensory feedback and internal predictions according to their reliability. Importantly, we further show that changes in movement speed led to urgency-dependent modulations of control gains. Our results highlight connections between motor control and decision-making processes, which both depend on the accumulation of sensory evidence and response urgency.

## Introduction

Studies on perceptual judgements and decision-making have suggested that the brain minimizes uncertainty by optimally combining cues across senses according to their respective reliability (Alais and Burr, 2004; Ernst and Banks, 2002; Van Beers et al., 1996). However, natural behavior often requires to make similar decisions during movement. For instance, when we reach for an object, the brain needs to decide how much to rely on vision and proprioception to guide the arm to the desired location. Thus, to understand the dynamics of multisensory perception and decision-making, it is important to investigate these processes during movement control.

Seminal studies on visual feedback responses during movement have documented how changes in reach end-points, and the modulation of the kinematics of corrective responses to visual perturbations, scaled with visual uncertainty and prior estimates (Izawa and Shadmehr, 2008; Körding and Wolpert, 2004; Tassinari et al., 2006). Izawa and Shadmehr (2008) interpreted their results as indicative of a continuously evolving estimate of the target state, as expected assuming the presence of a state-estimator similar to a Kalman filter. In more recent work, the same model has been suggested to explain the combination of proprioceptive and visual feedback in response to combined disturbances during human postural control and reaching tasks (Crevecoeur et al., 2016; Kasuga et al., 2022) but without considering the dependency of these responses on sensory uncertainty. Here we combined these two approaches to investigate whether visual uncertainty affects feedback responses to perturbation loads during reaching.

We performed two experiments in which we varied the temporal parameters of movement execution and of the visual feedback presentation to characterize the dynamical properties of the integration process. In Experiment 1, we instructed participants to perform fast and slow movements by varying the time allowed to reach the target. Thus, by changing movement time, we effectively varied the urgency to respond to the perturbation, which is known to be a determinant factor of decision-making and feedback control (Cisek et al., 2009; Poscente et al., 2021). In Experiment 2 we selectively increased the visual feedback presentation time during slow movements to assess whether this increased the contribution of vision as expected assuming an optimal observer (Kalman filter) that iteratively combines internal priors and sensory feedback over time. Such an evidence accumulation has also been shown to underlie deliberation during decision processes (Ratcliff, 1978).

Motor control theories based on stochastic optimal control (Todorov and Jordan, 2002) make clear predictions about the expected results. On the one hand, the control gains, with which the system responds to sensory errors, are known to be tuned to both the dynamics of the environment (Maurus et al., 2023) and the urgency to respond to a perturbation (Crevecoeur et al., 2013; Dimitriou et al., 2013; Oostwoud Wijdenes et al., 2011; Poscente et al., 2021) On the other hand, considering additive noise as a first approximation, the dynamics of state-estimation only depend on the statistics of the noise disturbances, and not on the urgency. Thus, in principle, there should be a transient effect of movement time on the modulation of feedback responses attributable to the control gains, while the estimation following visual errors should only depend on the sensory information available to the brain.

Our results were remarkably similar to the model predictions. First, assuming the presence of a Kalman filter could explain uncertainty-dependent modulations in feedback responses to multimodal perturbations. Second, simulating urgency-related modulation of feedback responses reproduced a transient increase in the visual feedback response for similar amounts of visual information that was observed in the data. Third, we observed that the contribution of visual feedback increased with viewing duration, which bared an obvious resemblance to a process of evidence accumulation over time. In all, we observed both an effect of urgency and evidence accumulation during multimodal perturbation responses, and suggest that these factors are separable computational operations underlying both movement control and decision-making.

## Materials and Methods

### Participants

This study is based on data collected from 32 healthy young adults aged 19 to 35. Sixteen (11 females) participants took part in experiment 1 and sixteen (10 females) in experiment 2. Handedness was assessed using the Edinburgh Handedness Inventory (Oldfield, 1971) and all participants reported to be righthanded. All participants had normal or corrected to normal vision and none of them indicated suffering from a neurological or motor disorder. Prior to participating in the experiment, participants were informed about the experimental procedure and gave written consent. All procedures were approved by the ethics committee at the host institution (*Comité d’Éthique Hospitalo-Facultaire*, UCLouvain). In total the experiment took three hours including information and preparation of the participant. All participants received a small financial compensation for their time.

### Apparatus and General Task Procedure

Both experiments were conducted using a KINARM Endpoint robotic device (KINARM, Kingston, Canada). The task paradigm was developed using Matlab’s Simulink and Stateflow toolboxes (Matlab 2015, Mathworks Inc. Natrick Ma, USA). During the experiment participants held the right handle of the KINARM robot and performed 20cm forward reaching movements with their right arm (Fig. 1a). The start and goal targets were displayed as gray circles with a radius of 1.2cm and were projected into the plane of the movement using a monitor-mirror setup. The start target was located 8cm from the bottom of the screen and 9cm to the right of the body midline and the goal target was located 20cm straight ahead from the start. Direct view of participants’ hands was blocked throughout the experiment but their hand position was indicated on the screen between trials using a 0.5cm-radius white cursor. During the movement, the visual feedback changed as explained in more detail below. To start each movement participants were instructed to move the cursor into the start target. Upon entry, the start target changed color from gray to blue and a waiting time interval of 2-4s (drawn from a uniform random distribution) was generated. The cue to initiate the movement was given by changing the color of the goal target to blue as well. In 75% of all trials a rightward constant load of 9N was applied to the hand as soon as it left the start target and remained on for the entirety of the trial (Fig. 1c). During the remaining 25% of trials participants experienced either no force or a leftward 9N force. The applied forces were ramped up and down over a period of 5ms.

**Figure 1.**
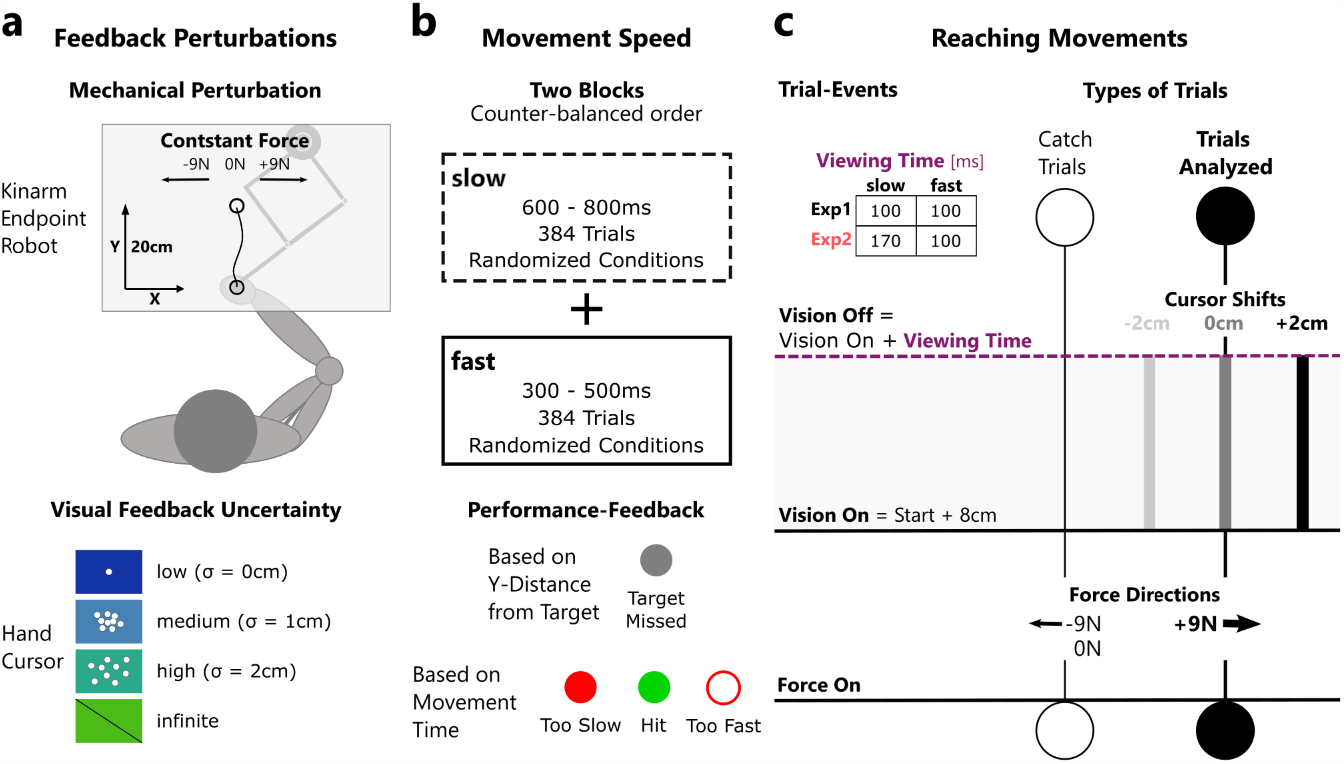
Experimental design. (a) Feedback Perturbations: Top panel: Participants performed 20cm right-arm reaching movements holding the handle of a robotic manipulandum. At movement onset a ±9N constant load or no load (0N) was applied to perturb the arm orthogonal to the movement direction. Bottom panel: Visual feedback was presented as a hand cursor and varied in uncertainty. In the condition of low uncertainty, a single hand cursor was presented (σ = 0cm), in conditions of medium and high uncertainty a cloud of 25 cursors was presented (σ = 1cm, σ = 2cm). In a fourth condition no visual feedback was presented (infinite). **(b) Movement Speed:** Top panel: The experiment consisted of two movement speed conditions that were applied in sessions of 384 trials each. The order of movement speed sessions was counter-balanced across participants. Movement speed was manipulated by imposing different timing constraints on participants’ movements. In the slow session, movements were counted as hits if the movement time was between 600-800ms and in the fast session between 300-500ms. Within each movement speed session, visual uncertainty conditions, force directions, and cursor shifts were applied in a random order. Bottom panel: Participants received feedback about the timing and length of their movement via a change in target color at the end of each movement. **(c) Reaching Movements:** Force perturbations were turned on when the movement was initiated and remained on during the movement and the stabilization phase. Visual feedback was presented for 100ms (Exp 1 fast & slow, Exp 2 fast) or 170ms (Exp 2 slow) once the hand crossed a distance of 8cm from the start target. Cursor shifts were only applied during trials with rightward 9N perturbations and the analyses focused on these trials only.

During the initial and final phases of the movement, no visual feedback of the hand position was shown to participants. However, when the hand crossed a threshold of 8cm straight ahead from the start target, visual feedback of the hand position was flashed on the screen for 100ms or 170ms (Fig. 1c; see details about exp. 1 and 2 below). To manipulate visual uncertainty, we presented either a single hand cursor, or a cloud of cursors of increasing spread, or no visual feedback about the hand location (Fig. 1a). We adapted this experimental manipulation from a previous study (Körding and Wolpert, 2004), but similar techniques have been used elsewhere (Ferrari & Noppeney, 2021; Tsay et al., 2021). In the single cursor condition the hand position was indicated by a white cursor of 0.3cm radius (low visual uncertainty). The cursor clouds were composed of 25 dots of 0.2cm radius. The position of each cursor was drawn randomly from a normal distribution centered on the hand position with a standard deviation of 1cm (medium uncertainty) or 2cm (high uncertainty). The trials without visual feedback served as a control condition with theoretically infinite visual uncertainty. On 50% of trials the center of the hand cursor or cursor cloud (even if invisible) was shifted 2cm to the left or right relative to the true location of the hand to induce discrepancies between the felt and seen hand location (cursor shift). This allowed us to quantify the influence of the visual feedback on the corrective response (see section Data Collection and Analyses for details). These cursor shifts were only imposed on trials with rightward force-perturbations to limit the total number of trials. Consequently, our analyses focused exclusively on those trials. Trials with leftward or no force perturbation served to make the task less predictable and keep participants focused.

We instructed participants to perform straight movements to the target and to stabilize their hand there for 2 seconds (stabilization phase). For trials with mechanical perturbations, they were asked to compensate for the force and stabilize their hand as close as possible to the goal target. For a successful trial, participants had to cross the distance of 20cm straight ahead from the start target within the imposed time interval (Fig. 1b; see details on Exp 1 below). However, trial success was not dependent on whether participants landed on target in the lateral dimension of the movement. Upon movement completion, participants received feedback in the form of a color-change of the goal target (Fig. 1b). There were four possible scenarios: if the goal target changed back to gray, it signaled that the participant had undershoot the target and that they should try to execute a longer movement on the next trial (these trials were excluded from further analyses); If the goal target filled red or turned to a red outline, it meant that the participant’s movement had been too slow or too fast respectively; Lastly, if the goal target changed to green, it indicated that the movement was performed correctly within the imposed time interval. To increase participants’ motivation, they gained one point for each successfully completed trial and were told to try to collect as many points as they could. Following the presentation of the feedback, the goal target was extinguished. To start the next movement participants had to move the handle back towards the start target and the hand cursor reappeared on the screen when the handle was within a 10cm radius of the start target. This was done to avoid that participants would notice shifts in the cursor position that were imposed during the experiment.

#### Experiment 1

The goal of experiment 1 was twofold. Firstly, we aimed to investigate the effect of visual hand feedback uncertainty on feedback corrections to a mechanical perturbation. Secondly, we aimed to study whether the response to the visual feedback additionally depended on the speed of the performed movement. During this experiment visual feedback was presented for 100ms during the movement across two different conditions of movement speed (Fig. 1c).

The experiment was divided into two separate parts, one in which the movement time window was set between 300ms and 500ms (fast movement speed condition), and a second condition in which movement time was between 600ms to 800ms (slow condition; Fig. 1b). In each condition, movements were rewarded with a point if they were performed within the instructed time intervals. Each part was preceded by two practice blocks of ten movements without force perturbation, followed by twenty movements with load perturbations (ten leftward and ten rightward), so that participants could familiarize themselves with the task and the movement speed requirement. If they failed more than 50% of the practice movements, the training block was repeated once. The order of speed conditions was counterbalanced across participants to limit potential biases linked to the first practiced condition.

Each condition was divided into 6 blocks of 64 trials each. Each of these blocks consisted of 48 trials with a rightward force perturbation (4 repetitions of 4 visual uncertainty by 3 cursor shifts combinations), 8 trials with leftward force perturbation, and 8 trials without force (2 repetitions of 4 possible visual uncertainty conditions). The order in which these trials appeared was completely randomized. When needed, participants were allowed to take short breaks between blocks. Usually a longer break of about 5 minutes was inserted between the first and second part of the experiment. In total each participant performed 768 trials and the experiment lasts about 2.5h in total. At the end of the experiment, each participant was asked whether they noticed something during the experiment, but none of them reported having noticed the cursor shifts.

#### Experiment 2

Experiment 2 was performed as a follow-up experiment to investigate whether the differences between fast and slow movements observed in experiment 1 could be at least partly explained by the difference in distance during which the visual feedback was visible. Put differently, we were interested to see whether increasing the duration of the visual feedback during slow movements would increase the visual contribution to movement corrections. In experiment 1 we imposed a fixed 100ms time interval during which the visual feedback was shown. Given the different movement speeds, this resulted in a difference of movement distance during which visual feedback was available in the fast and slow conditions. Analyses revealed, that in experiment 1 visual feedback was present on average (mean ± SD) for 5.7cm (± 0.9cm) in the fast condition and for 3.2cm (± 0.7cm) in the slow condition. To account for the potential influence of this difference on our result of Experiment 1, we designed a second experiment in which we matched the distance travelled with visual feedback between the fast and slow movement condition based on the difference in group average velocity. To achieve this, we extended the visual feedback viewing time in the slow condition to 170ms (Fig. 1c). This resulted in an average distance travelled with visual feedback of 5.6cm (± 0.9cm) in the fast and 5.3cm (± 1.2cm) in the slow condition. Importantly, the distance at which the visual feedback was turned on remained at 8cm from the start target in both movement speed conditions. All other experimental procedures also remained identical to Experiment 1.

### Kinematics: Data collection & analyses

During the experiment we recorded kinematic data at a sampling rate of 1kHz using KINARMS’s Dexterit-E software (Version 3.9). Preprocessing of the data was performed using custom written MATLAB scripts (MATLAB 2021a, Mathworks Inc. Natrick Ma, USA). The recorded positions and forces were filtered using a low-pass, fourth-order, dual-pass Butterworth filter with a cutoff frequency of 50Hz. Hand velocities and accelerations were derived from the raw position data using a fourth-order central difference approximation and passed through the same filter.

To quantify the influence of the visual feedback on movement corrections, we extracted the lateral hand position during rightward perturbation trials at different time points during the movement following the onset of the visual feedback. In particular, we extracted this information for each participant at 150ms, 300ms, 500ms, 700ms, and 900ms following the onset of visual hand feedback. It should be noted that each of these timepoints lay outside of the 100ms time window during which vision was presented in Experiment 1. We chose these specific timepoints based on visual inspection of the data as they track well the development of the influence of the visual feedback. Next, we computed the regression slope between the cursor shift that was applied and the lateral position of the hand relative to the center of the target at each timepoint. This regression slope served as a quantification of how much participants relied on the visual feedback. A slope of 0 signified that there was no shift in the lateral position of the movement relative to the cursor shift, and hence no influence of the visual feedback. On the contrary, a slope of -1 meant that participants fully compensated for the 2cm lateral cursor shift. Lastly, we computed the average regression slopes across all participants and compared these values across the conditions of visual uncertainties and movement speeds.

Additionally, we computed the variability of positions at the same five timepoints during the movement (150ms, 300ms, 500ms, 700ms, and 900ms) based on the 2-dimensional dispersion ellipses of X- and Y-positions for each visual uncertainty condition using singular value decomposition and defined the variability as the area of the ellipse. Finally, we looked at the lateral forces applied to compensate for the perturbation. For this analysis, we first computed the difference in average lateral forces between the rightward or leftward cursor shift condition and the no cursor shift condition for each participant separately. Next, we computed the difference of this difference ((rightward cursor shift - no cursor shift) - (leftward cursor shift - no cursor shift) and extracted the maximum value of this quantity.

### Electromyography (EMG): Data collection & analysis

We recorded EMG (electromyography) signals from the pectoralis major (PM) and posterior deltoid (PD) muscles in the right shoulder. These muscles act as agonists to the right- and leftward force perturbation, respectively. Muscle activity was recorded using bipolar surface electrodes (DE-2.1. EMG Sensor, Delsys) which were attached over the muscle belly. Prior to applying each electrode, we cleaned the skin underneath with cotton gauze and medical alcohol and coated the contacts of each electrode with conductive gel to enhance the signal-to-noise ratio. Depending on the signal strength of each participant, we amplified the signal by a factor of 1,000 or 10,000 (Bagnoli-8 EMG System, Delsys). All EMGs were recorded at a sampling frequency of 1kHz.

The preprocessing and analysis of the muscle recordings was performed using custom-written MATLAB scripts (MATLAB 2021a, Matworks Inc. Natrick Ma, USA). First, we aligned the EMG recordings to movement onset and bandpass filtered the signal using an eighth-order, dual-pass Butterworth filter (cut off frequencies: [20, 250] Hz). After filtering, signals were rectified and normalized by the average activity computed based on 4 separate normalization blocks which were performed before and after each speed-condition of the experiment. During these calibration trials, participants were presented a 2 by 2cm square on the screen in front of them. As soon as they moved their hand inside the square a 9N constant force was applied to the left or right against their hand to activate one of the two muscles of interest. This force remained on for 2s and participants were instructed to counter the force and keep their hand inside the square on the screen. For the normalization, we extracted a 1s recording between 0.5s and 1.5s following the onset of the force. Next, we computed the mean rectified muscle activity across this time window for all the repetitions of each force-direction. Finally, the activity measured in each muscle during all trials was divided by their corresponding calibration values.

To investigate the influence of visual uncertainty on the modulation of muscle responses, we realigned the preprocessed EMG recordings to the presentation onset of the visual feedback and computed average traces for each visual uncertainty and cursor shift condition across participant. To improve the illustration of the group-average EMG traces, we plotted a moving average with a window size of 11 samples. Next, we computed the difference between conditions with a cursor shift to the right and left to illustrate the effect of the visual uncertainty on the modulation of muscle activity with the direction of the cursor shift. For illustration of these delta EMG traces, we again plotted a moving average with a window size of 31 samples across 0 – 300ms following the onset of the visual feedback. Lastly, to compare the different visual uncertainty conditions statistically, we computed the average delta EMG responses over a time window from 100 – 250ms following the onset of the visual feedback.

### Experimental Design and Statistical Analyses

We performed our statistical analyses using custom written MATLAB scripts (MATLAB 2021a, Matworks Inc. Natrick Ma, USA). For our main analyses, we used a four by two repeated measures ANOVA with visual uncertainty and movement speed as within participant factors. We assessed the main effect of visual uncertainty and movement speed as well as their interaction on the slope and the movement variability at each of the five specified time points following the onset of the visual feedback (150ms, 300ms, 500ms, 700ms, and 900ms). Further, we analyzed the influence of visual uncertainty and movement speed on the absolute and delta lateral forces as well as the delta EMG. Lastly, we compared the slopes across experiments 1 and 2 using a repeated measures ANOVA with movement speed as within participant factor and experiment number as between participant factor. For each ANOVA we report the F-statistic, the degrees of freedom, the P-value and the partial eta squared as a measure of effect size (Lakens, 2013). To highlight significant mean differences in our figures, we computed post-hoc pairwise comparisons with Bonferroni corrections. For within-participant comparisons all statistical results with a corrected P-value < 0.005 are considered significant (Benjamin et al., 2018; Lakens et al., 2018). Additionally, results with a P-value < 0.05 are interpreted as a significant trend. For between-participants comparisons, statistical results are regarded as significant if the P-value was below a threshold of 0.05.

### Model

We used an optimal control model to simulate the influence of visual uncertainty and movement speed on feedback corrections to combined mechanical and visual perturbations. Our model describes the translation of a point mass (*m* = 1kg) in a plane. Such a simplified linear model has been previously used to approximate the nonlinear behavior of a multi-jointed arm (Izawa and Shadmehr, 2008; Nashed et al., 2012). The model includes a damping factor *G* = 0.1 Nsm^-1^ and we approximated the muscle dynamics using a first-order low-pass filter with time constant *τ* = 66ms (Brown and Loeb, 2000). The state variables include the hand position (*x*) and velocity 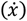, the commanded force (*F*_*x*_) produced by the control input (*u*), and the external force used to simulate the mechanical perturbation (*F*_*E*_). Additionally, we added a variable for the cursor motion (*x*_*c*_), as well as a variable to define an offset between the cursor and hand motion (*x*_*off*_). This procedure was chosen to allow dissociating the hand from the cursor as in the experiment (cursor shift), and make the offset variable non-observable such that any shift between cursor and hand had to be estimated (these modeling choices are not the unique way of dissociating hand and cursor position). Thus, the state vector is defined as follows:

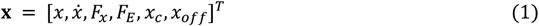

Please note, that we differentiate between the state vector **x**, in bold font, and the position variable *x*, in italicized font, which is an entry in the state vector.

Finally, we augmented the state vector with the target state variables. The continuous differential equations of the system are the same for *x* and *y* dimensions without interaction. For simplicity we only describe the dynamics in the *x* dimension here corresponding to the lateral hand coordinate:

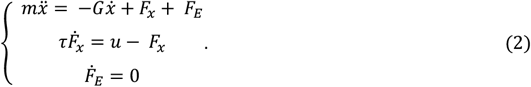

The third equation expresses that changes in the external force input are assumed to follow a step-function.

Next, the dynamics of the system were discretized using Euler integration with a time step of *δt* = 10ms. This led to the following representation of the discrete-time control system:

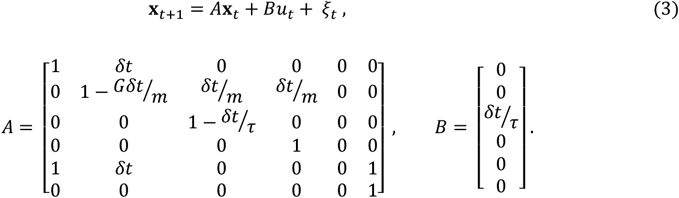

*ξ*_*t*_ is an additive multivariate Gaussian noise with zero-mean and known covariance matrix. The last two rows of the matrices A and B correspond to the cursor position and the offset between cursor and hand, respectively. Calling *x*_*c,t*_ the cursor position and *x*_*off,t*_ the offset between hand and cursor at time *t*, the discrete time dynamics of these variables are respectively *x*_*c,t*+1_ = *x*_*t*+1_ + *x*_*off,t*_ and *x*_*off,t*+1_ = *x*_*off,t*_.

As mentioned above, we assumed observability of all state variables except the offset between hand and cursor position. Hence, the observation matrix *H* is defined as *diag*(1,1,1,1,1,0) and the feedback equation can be written as follows:

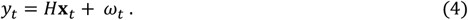

*ω*_*t*_ is the sensory noise with covariance matrix *Σ*_*ω*_. We manipulated the visual uncertainty by increasing or decreasing the corresponding element in *ω*_*t*_ to simulate different amounts of noise in the feedback about the cursor position. This was done arbitrarily by multiplying or dividing *Σ*_*ω*_ by a factor of 10. This procedure was constrained by the experimental design in which the sensory signal was actually increased or decreased, but the factors used could not match noise statistics of visual estimates accurately, so we verified that they produced differences in slopes that were broadly consistent with the behavioral observations.

Next, we used Kalman filtering to obtain maximum-likelihood estimates of the system state at each timepoint. This estimator assumes an optimal combination of prior and sensory feedback which relies on internal knowledge of state-space representation matrices (*A, B* and *H*), control input, and noise covariance matrices. The prior is the expected value of the next state given the current estimate 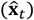 and the control input (*u*_*t*_) and is computed by simulating the system dynamics over one timestep:

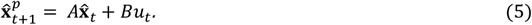

The estimated state at the next timestep is then computed by combining the prior with the observed feedback error weighted by the Kalman gain (*K*):

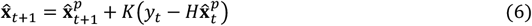

Please note, to ensure corrective responses to the mechanical and visual perturbations, Kalman gains were computed using non-zero noise for the external force and the hand-cursor offset. This allows the estimator to infer jumps in this variable, which is otherwise impossible if it is assumed that there is no noise affecting these variables. The actual noise in the simulations for these variables was then set back to zero in agreement with their physical properties.

We computed optimal control gains using a quadratic cost function with a penalty on position error and control:

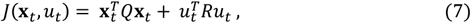

where *R* = 10^-4^ describes the cost to penalize large control commands and Q represents the cost applied to position errors during the 500ms stabilization phase at the end of the simulated movements. Thus, Q was zero throughout the movement and applied a quadratic cost term to the difference between cursor and target position during the stabilization phase. The resultant optimal control policy is a linear function of the estimated state:

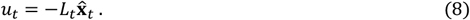

with *L* representing the optimal control or feedback gains. Details about the derivation of the controller can be found elsewhere and followed standard techniques (Åström, 1970; Todorov, 2005).

To simulate mechanical perturbations, we set the external force (*F*_*E*_) to 9N during a simulation run as soon as the *y* position of the state exceeded a distance of 0.5cm from the start position. Additionally, we added a shift of the cursor at 8cm from the start position by setting the offset between hand and cursor position (*x*_*off*_) to ±2cm. We computed 25 simulation runs with movement time 400ms and 700ms for fast and slow movements, respectively. At the end of each movement we added a 500ms stabilization phase to mimic the experimental paradigm as closely as possible.

## Results

One of the hallmarks of optimal multisensory integration is that the contribution of each sensory cue is weighted by the inverse of its variance (reliability). Therefore, Experiment 1 aimed to investigate whether visual uncertainty also modulated online feedback responses to mechanical loads applied during reaching. Moreover, we varied movement time to test whether this influences the dynamics of corrective responses to combined force and visual perturbations.

Figure 2a displays the average trajectories during rightward perturbations for one exemplar participant in Experiment 1. The different directions of the cursor shift are shown in different shades of gray. Similarly, panel b shows the average lateral positions for the same participant. The dashed vertical lines indicate the median onset and offset time of the visual feedback for this participant. In trials in which visual feedback was presented during the movement, we can observe a clear divergence of the lateral positions in correspondence with the direction of the cursor shift. Specifically, when the cursor was shifted 2cm to the right relative to the hand coordinate, the participant’s corrective response increased. Conversely, when the cursor was shifted 2cm to the left, closer to the midline of the movement, the correction was reduced. Thus, the visual feedback clearly influenced the feedback response to the force perturbation.

**Figure 2.**
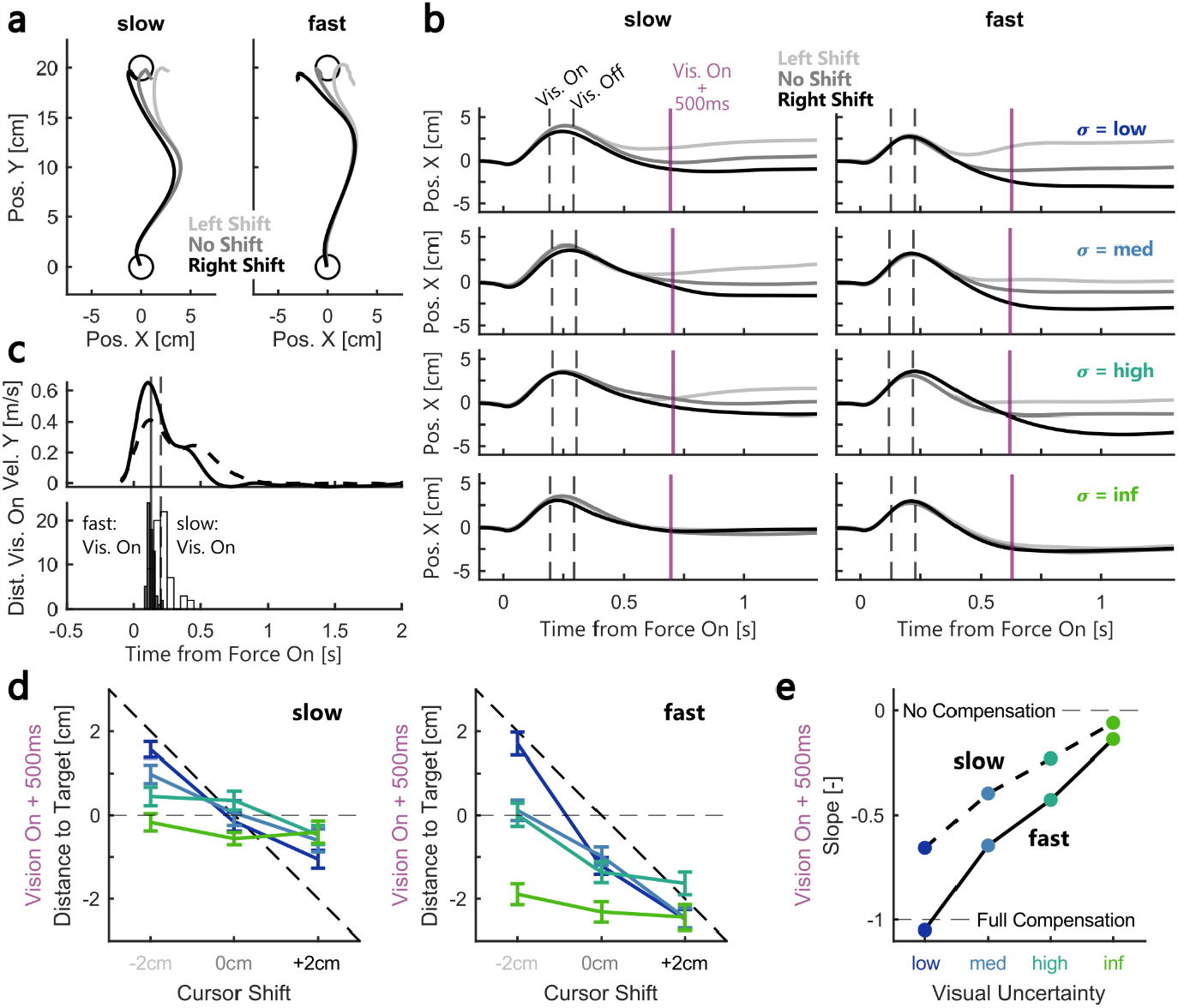
Example participant behavior. **(a)** Average hand trajectories during low visual uncertainty for one example participant in experiment 1. Trajectories during slow movements are shown on the left panel, those during fast movement on the right panel. Shades of grey represent the three cursor shift conditions. **(b)** Average lateral position over time for the same example participant. The left column represents averages across slow movements (visual uncertainty low – inf are shown in top to bottom panel). The right column represents averages across fast movements. The dashed lines represent the onset and offset of the visual feedback and the purple line marks the timepoint 500ms after vision onset. **(c)** Top: Average forward velocities during fast (full line) and slow (dashed line) movements without cursor shift and with low visual uncertainty for the same example participant. Bottom: Histograms of the onset times of the visual feedback relative to force onset for the same participant. The slow condition is shown as open bars and the fast as filled bars. Vertical lines represent the median onset time during slow (dashed) and fast (full) conditions. **(d)** Left: Average lateral positions extracted 500ms after vision onset relative to the target center as a function of the cursor shift during slow movements. Blues and greens represent the different visual uncertainties. Right: Same as left but for fast movements. **(e)** Average slopes between cursor shift and lateral hand position as a function of visual uncertainty and movement speed (slow = dashed, fast = full) for the same example participant.

The top panel of Figure 2c shows the average forward velocity during fast (full line) and slow (dashed line) movements for the same example participant, while the bottom panel depicts the distribution of the onset times of the visual feedback. The vertical lines indicate the median onset time in each movement speed condition for this participant. It is visible that the onset of the visual feedback occurred after peak velocity in both conditions.

Next, to compare the contribution of the visual feedback across different levels of uncertainty and movement speed, we extracted the lateral positions at 150ms, 300ms, 500ms, 700ms, and 900ms after the onset of the visual feedback. Figure 2d shows the lateral positions at 500ms as a function of the cursor shift for the same exemplar participant during slow (left panel) and fast (right panel) movements. We computed the slope across the cursor shifts and plotted these values as a function of the visual uncertainty (Fig. 2e). A slope of 0 indicates that the movement corrections did not differ depending on the cursor shift, whereas a slope of -1 signifies that the participant compensated fully for the shift of the cursor or the expected value of the cursor-cloud. Intermediate values of the slope indicate a partial correction for the visual shift. Figure 2e clearly illustrates that the contribution of the visual feedback to the movement correction decreased with visual uncertainty in both movement speed conditions.

To quantify the effect of visual uncertainty and movement speed on feedback corrections at the group level, we computed the slopes for all participants at the five selected time points following the onset of the visual feedback presentation. Figure 3a shows the average hand position at each of the five timepoints during the movement and stabilization phase. Naturally, these positions differed between the slow and fast condition due to the difference in movement speed. From 500ms following the presentation of the visual feedback the hand stabilized close to the target for both slow and fast conditions. Figure 3b demonstrates that 150ms after the onset of the visual feedback, there was no observable influence of vision on the movement correction. However, starting at 300ms, fast movements began to show negative slopes, signifying a modulation in lateral hand position in accordance with the shifted cursor. Even later during the movement, at 500ms following the onset of vision, we observed negative slopes that scaled with visual uncertainty in both movement speed conditions, but the slopes were clearly larger during fast movements. Importantly, this difference between slow and fast conditions decreased again during even later timepoints (700ms and 900ms). We tested these effects using a repeated measures ANOVA with visual uncertainty and movement speed as within participant factors which revealed a main effect of visual uncertainty starting at 300ms after vision onset 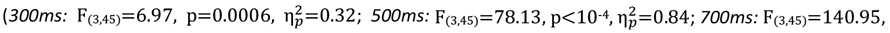 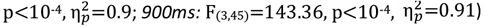. In addition, slopes at 300ms and 500ms where significantly larger during fast compared to slow movements 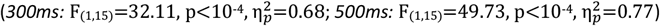. At 700ms the relationship between fast and slow slopes showed a clear significant trend 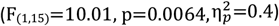 but the effect was not significant anymore at 900ms (F_(1,15)_=2.41, p=0.14). Taken together, these results highlight that feedback responses decreased with visual uncertainty. Additionally, there was a transitory period between 300-700ms following the onset of vision during which the slopes were larger during fast movements.

**Figure 3.**
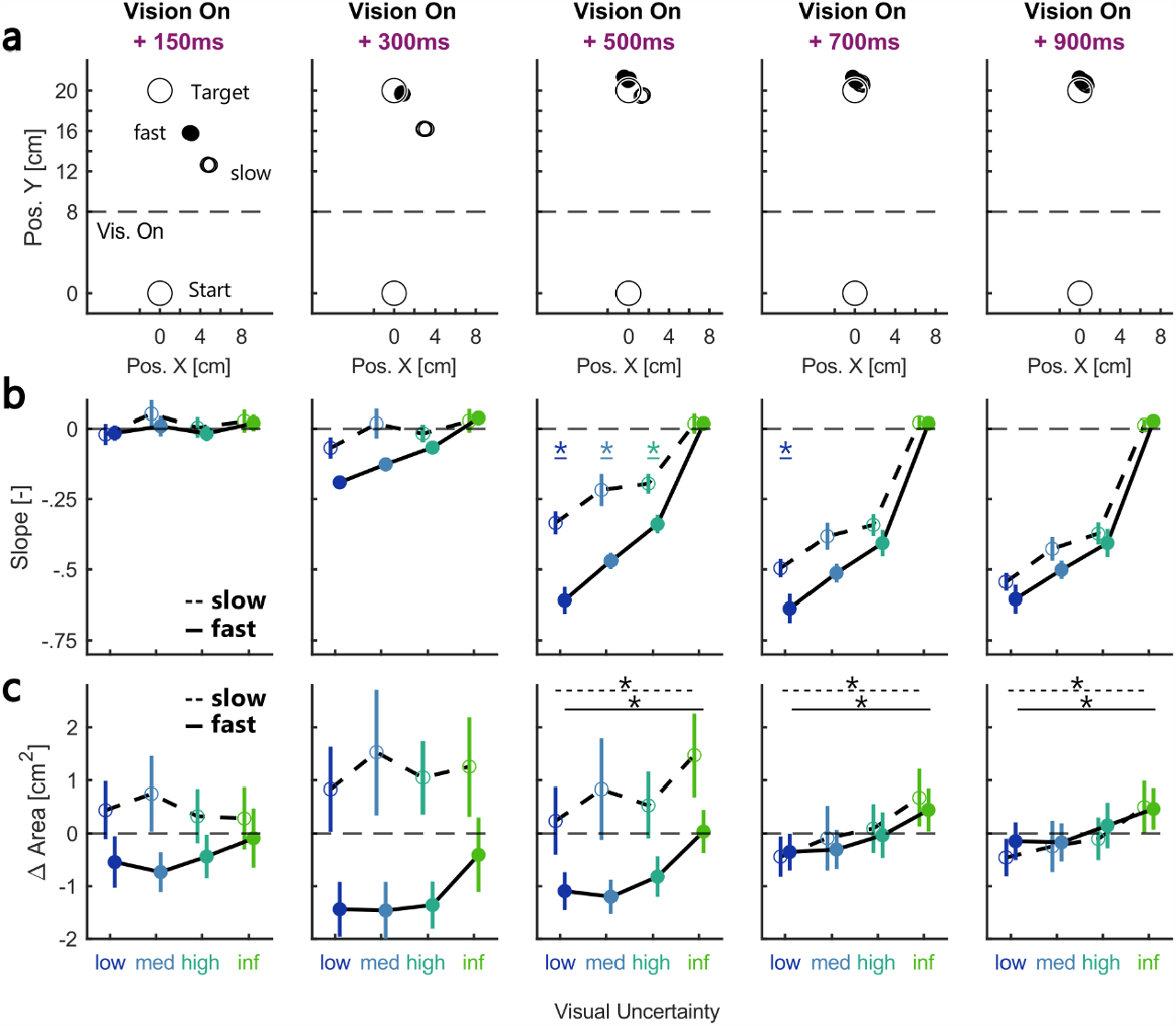
Influence of visual feedback over time. **(a)** Group average location of the hand relative to the target during the two movement speed sessions at different timepoints relative to vision onset. Hand positions were averaged across trials without cursor shift and open circles represent slow and filled circles fast movements, respectively. **(b)** Group average slopes computed at different timepoints after the onset of the visual feedback as a function of visual uncertainty. The slopes were averaged across all participants in experiment 1. **(c)** Group average variability in hand positions over time as a function of visual uncertainty. Variability was computed as the area of the ellipse described by the SD of x- and y-positions. Each panel represents the change in area relative to the total average at that timepoint. The color-coding is identical to the one used in figure 2 and the error-bars indicate group-averages ± SEM. * indicate pairwise comparisons with p < 0.005. In panels (c) lines on top of the panels indicate the pairwise comparison between conditions with low and infinite visual uncertainty in the slow (dashed line) and fast (full line) conditions.

If visual information was indeed integrated in an optimal manner, movement variability should increase with visual uncertainty. Figure 3c shows the average area of the ellipse describing variability in X- and Y-positions. For illustrative purposes these variability values are shown as the difference to the average across all uncertainty conditions at each timepoint. We can observe that the variability in positions was larger during slow movements compared to fast up until 700ms following the onset of visual feedback. The larger variability during slow movements might be linked to reduced temporal alignment of the traces, which is also visible in the wider distribution of the visual feedback onset times (Fig. 2c). Importantly, starting at 500ms, we can see a positive relationship between the level of visual uncertainty and the variability (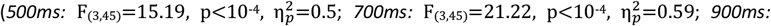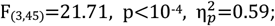; repeated measures ANOVA). This further supports that the visual feedback contribution to the movement correction scaled with the signal reliability.

In Experiment 1 we presented the visual feedback for 100ms during slow and fast movements, consequently, visual information was available for a larger fraction of the hand path during fast movements. Hence, we performed a second experiment during which we matched the distance of the movement travelled with visual feedback between slow and fast movements. We used the average velocity to estimate the additional viewing time necessary to match the observable travelled distance, and accordingly, increased the visual feedback presentation from 100ms to 170ms during slow movements in Experiment 2 (Fig. 4a top & bottom panel; see Methods). Importantly, the average peak forward velocities remained identical between Experiment 1 and 2 (Fig. 4b top & bottom panel; Exp. 1 (mean ± SD): slow: 0.4m/s ± 0.07m/s, fast: 0.65m/s ± 0.1m/s; Exp. 2: slow: 0.4m/s ± 0.07m/s, fast: 0.64m/s ± 0.1m/s). At 500ms, 700ms, and 900ms after vision onset, we observed slightly larger slopes during the slow movements in Experiment 2, while there was no such difference between experiments in the fast movements (Fig. 4c & d). Statistically, we observed an interaction between the effect of movement velocity and the experiment at 700ms and 900ms (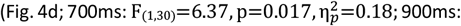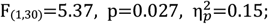; repeated measures ANOVA). In particular, while the slopes were significantly larger in the fast compared to the slow condition in Experiment 1, this difference was absent in Experiment 2. Hence, increasing the viewing duration of the visual feedback during slower movements increased the corrections for the cursor shift.

**Figure 4.**
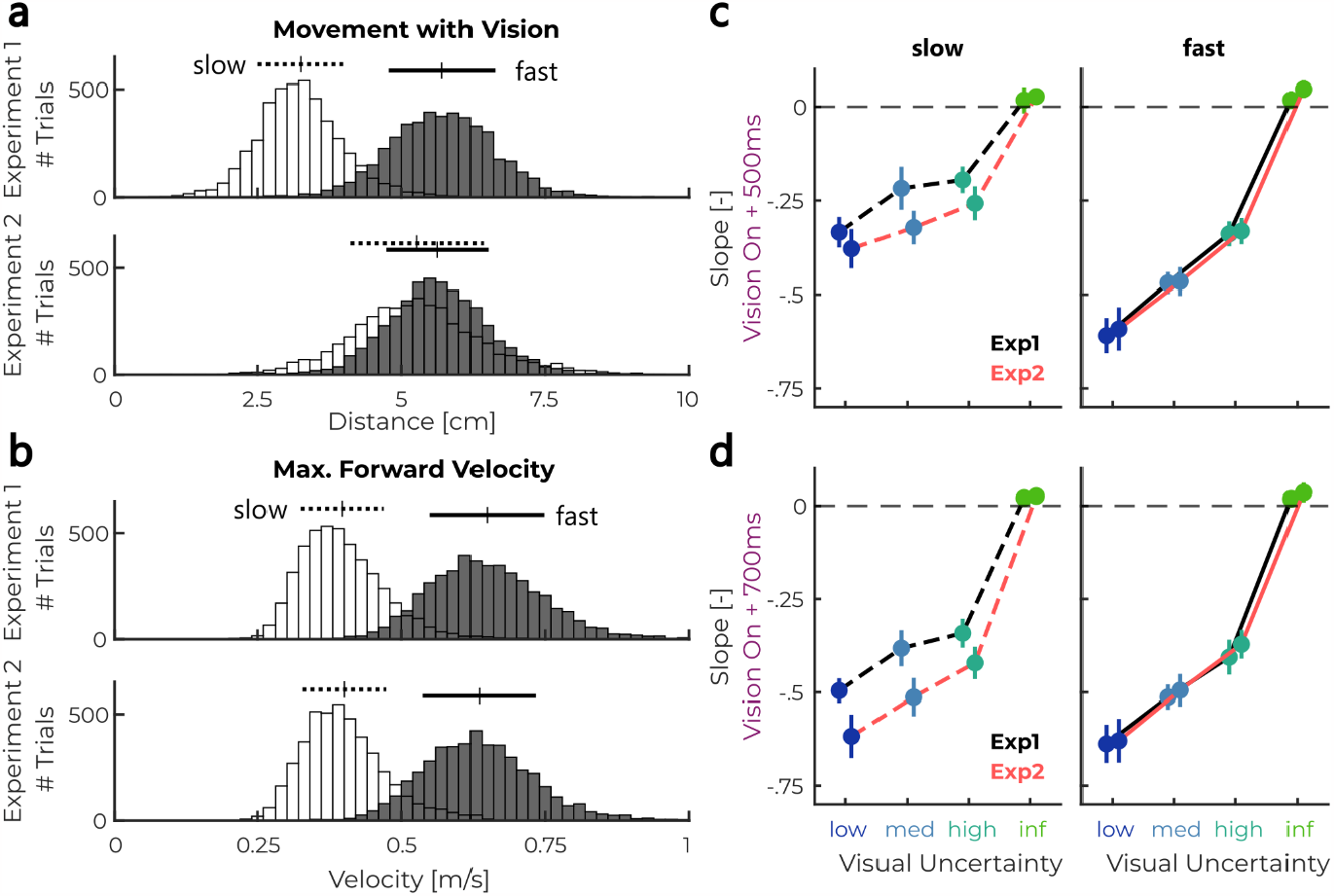
Influence of presentation time of the visual feedback. **(a)** Distribution of movement distances performed in the presence of visual feedback across all participants in experiment 1 (top) and experiment 2 (bottom). Slow movements are shown in white and fast movements in black. Bars on the top of the histograms indicate mean values ± 1SD. **(b)** Distribution of maximum forward velocities across all participants in experiment 1 (top) and experiment 2 (bottom). Bars on the top of the histograms indicate mean values ± 1SD. **(c)** Slopes computed at 500ms after the onset of the visual feedback as a function of visual uncertainty during slow (left) and fast (right) movements. Data from experiment 1 is shown in black and data of experiment 2 in red. **(d)** The same as (c) but at 700ms after the onset of the visual feedback. The color-coding is identical to the one used in figure 2 and the error-bars indicate group-averages ± SEM.

Previous studies have shown that control gains increase with the urgency to respond to a perturbation (Crevecoeur et al., 2013; Poscente et al., 2021). To assess whether this was the reason why we observed stronger responses to the visual feedback during fast movements, we compared the lateral forces applied by participants during trials with a rightward perturbation load across movement speed conditions (Fig. 5a, data Exp. 1). The maximum absolute lateral forces were clearly larger during fast compared to slow movements (Fig. 5b; Exp. 1: 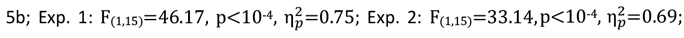; repeated measures ANOVA), which demonstrates an overall increase in responses to the force perturbation. We observed no influence of the cursor shift or the visual uncertainty on the peak lateral forces. To further investigate the influence of visual feedback uncertainty on the force responses, we computed the difference in forces applied during right and left cursor shifts (delta force; Fig. 5c). Next, we extracted the maximum delta force for each visual uncertainty condition (Fig. 5d). Using a repeated measures ANOVA, we observed a main effect of movement velocity (Exp. 1: 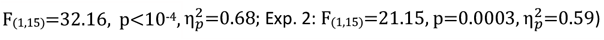) as well as a main effect of visual uncertainty (Exp. 1: 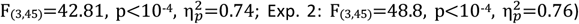) on the maximum delta force. Additionally, there was a significant interaction between movement velocity and visual uncertainty on the maximum delta force in Experiment 2 (Exp. 1: 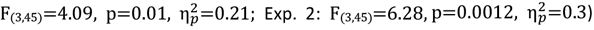). These findings demonstrate that the larger contribution of vision during fast movements observed in the slopes shown in Figure 3, could be linked to an increase in control gains with movement speed.

**Figure 5.**
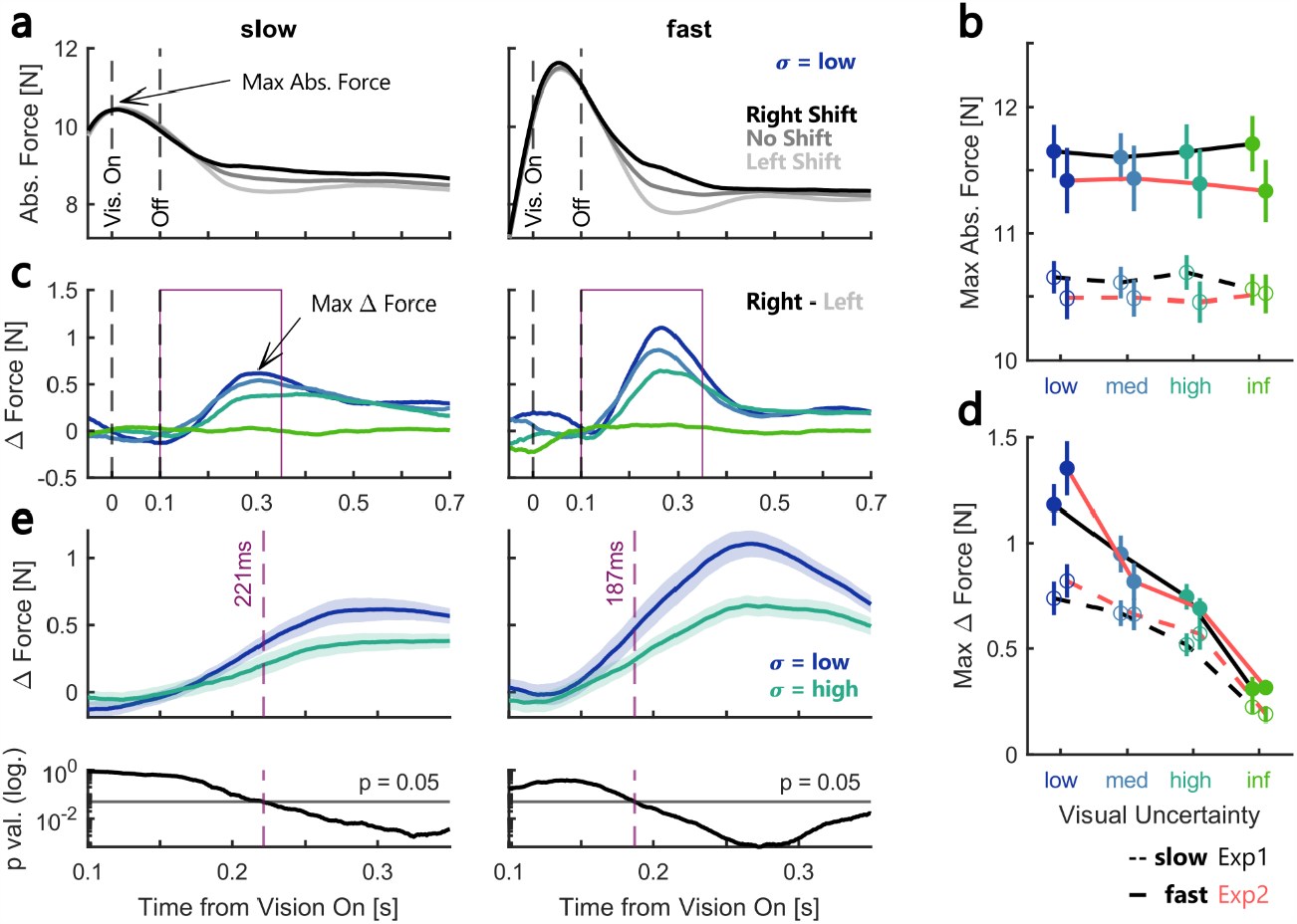
Lateral forces scale with movement speed and visual uncertainty. **(a)** Group average absolute lateral force during rightward force perturbations and low visual uncertainty in the slow (left) and fast (right) movement condition in experiment 1. Data was aligned to the onset time of the visual feedback. Shades of gray correspond to the three different cursor shift conditions. **(b)** The maximum lateral force indicated in panel (a) as a function of visual uncertainty. The full line represents fast and the dashed line slow movements. Data of experiment 1 are shown in black and experiment 2 are shown in red. **(c)** Difference in lateral forces to rightward - leftward cursor shift during slow (left) and fast (right) movements in experiment 1. **(d)** The maximum delta force indicated in panel (c) as a function of visual uncertainty. **(e)** Top: Zoom-in of the delta lateral forces for conditions with low (blue) and high (turquoise) visual uncertainty during slow (left) and fast (right) movements. The area that is zoomed-in is depicted by a purple box in panels (c). The shaded area corresponds to mean ± SEM. Dashed purple lines mark the timepoint when the two traces started to diverge (determined by a running t-test). Bottom: P-values of running t-tests over time. Dashed purple lines mark the moment the p-values falls below 0.05. The color-coding is identical to the one used in figure 2 and the error-bars indicate group-averages ± SEM.

To determine the onset of the effect of visual uncertainty on the force differences, we computed running t-tests to see when the force traces started to differ between the conditions with low and high visual uncertainty. We defined the onset as the first timestep after the removal of the visual feedback at which the P-value crossed below a threshold of 0.05. The estimated onset of the difference was 221ms during slow movements and 187ms during fast movements in Experiment 1. Further, we used bootstrapping with 10,000 iterations to estimate distributions of onset times for the slow and fast movements and observed no significant difference in onset times between movement speed conditions (data not shown). Thus, independent of movement speed, uncertainty dependent responses to the visual feedback occurred at ∼200ms following the onset of vision.

The increase in feedback responses with movement speed was also visible in the muscle activity recorded in the pectoralis deltoid muscle of the right shoulder (Fig. 6a (Exp. 1) & 6c (Exp. 2)). To investigate the influence of visual uncertainty on the modulation of muscle responses, we computed the average EMGs for each combination of cursor shift and visual uncertainty. The top panels of Fig. 6a and 6c show the group-average for each cursor shift during the condition with low visual uncertainty. For both slow and fast movements, we can see a separation of the traces depending on the cursor shift direction at around 100ms following the onset of the visual feedback. Next, to determine whether this effect was modulated by visual uncertainty, we computed the difference in EMGs between the right and left cursor shift for each visual uncertainty condition. The lower panels in Fig. 6a and 6c illustrate that this difference clearly decreased with increasing uncertainty. We then computed the average delta EMG across a time window ranging from 100 to 250ms following the onset of the visual feedback (Fig. 6b (Exp. 1) & 6d (Exp. 2)). The average activity was overall larger during fast movements (Exp. 1: 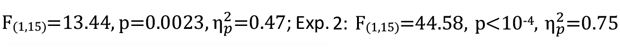; repeated measures ANOVA) and showed a clear scaling with visual uncertainty (Exp. 1: 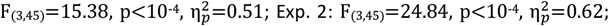; repeated measures ANOVA).

**Figure 6.**
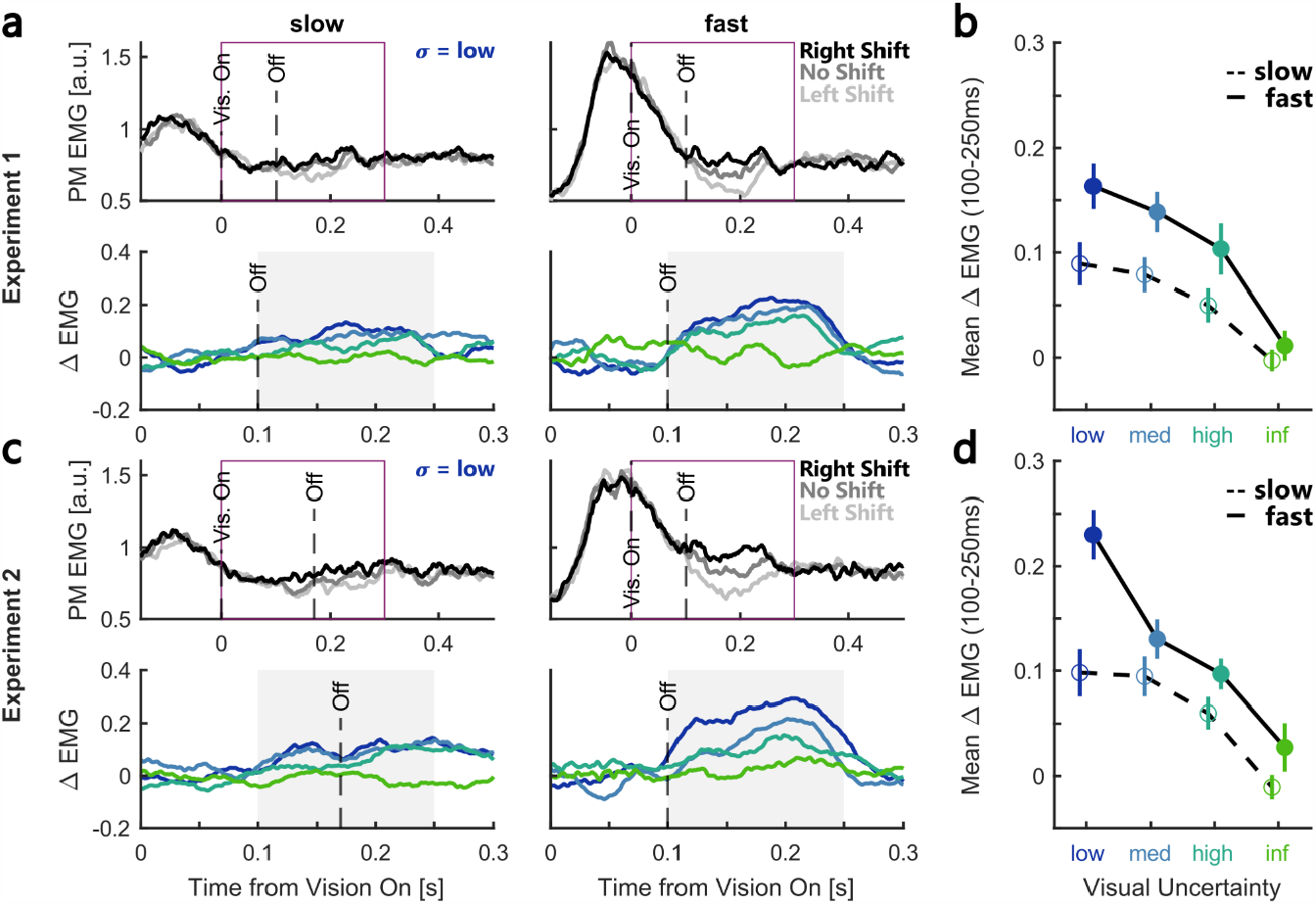
Muscle responses scale with movement speed and visual uncertainty. **(a)** Top: Group average pectoralis major EMGs during rightward force perturbations and low visual uncertainty in the slow (left) and fast (right) movement condition in experiment 1. Data was aligned to the onset time of the visual feedback. Shades of gray correspond to the three different cursor shift conditions. Dashed black lines indicate onset and offset of visual feedback. Bottom: Zoom-in of the difference in EMGs to rightward - leftward cursor shift during slow (left) and fast (right) movements in experiment 1. The area that is zoomed-in is depicted by a purple box in the top panels. **(b)** EMG responses averaged across 100-200ms following vision onset (gray-shaded area shown in (a) bottom panels) as a function of visual uncertainty. The slow condition is represented by a dashed line and open circles, the fast condition by a full line and filled circles. **(c)** Same as (a) but for experiment 2. **(d)** Same as (b) but for experiment 2. The color-coding is identical to the one used in figure 2 and the error-bars indicate group-averages ± SEM.

We implemented an LQG controller with a state estimator based on a Kalman filter to test whether such a continuous integration model could capture the observed effects of visual uncertainty and movement speed on feedback corrections. As in our experiments, a 9N constant load was applied to the simulated point-mass as soon as it left the start position. For simplicity, contrary to the experiment, visual feedback was present during the entire length of the simulated movements as it is difficult to model a transient presentation of visual information in a linear system. Instead of the sudden flashing of the visual feedback, we introduced an instantaneous right-or leftward cursor jump at the moment when the visual feedback was presented in the experiments. This cursor jump was not directly observable through the feedback equation and had to be estimated by the state-estimator. In spite of all simplifying assumptions, the simulated trajectories closely resembled the observed behavior and displayed a similar divergence in lateral hand positions based on the direction of the cursor jump (Fig. 7a). Importantly, the model shows that the rate at which the state estimation error decreases following the cursor jump depends on visual uncertainty but does not differ between movement speeds (Fig 7b). As shown in Figure 7c, the model accurately predicted an increase in the force modulation with visual feedback during fast movements, which was the result of a time-dependent increase in control gains.

**Figure 7.**
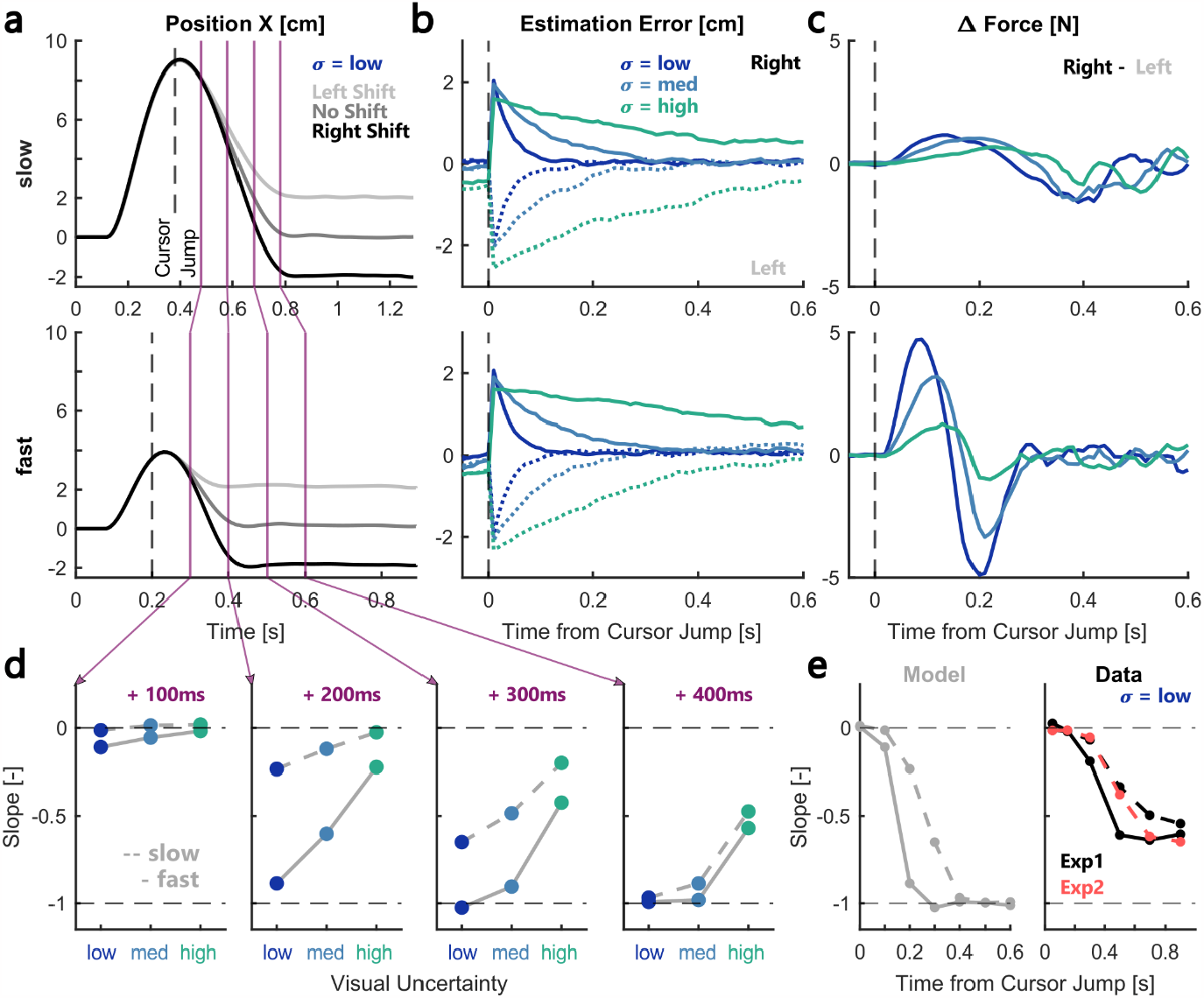
Model results. **(a)** Simulated lateral positions over time during slow (top) and fast (bottom) movements with low visual uncertainty. The dashed line marks the moment the cursor jump was applied and the shades of gray correspond to the direction of the jump. Purple vertical lines correspond to 100, 200, 300, and 400ms after the cursor jump. **(b)** Average decay of the estimation error following the cursor jump during different visual uncertainties (blues & greens). The decay following a leftward cursor jump is shown as dotted lines, that following rightward jumps as full lines. **(c)** Difference in lateral forces to rightward - leftward cursor jumps during slow (top) and fast (bottom) simulated movements. **(d)** Slopes of the simulated movements at the four highlighted timepoints following the cursor jump as a function of visual uncertainty. Slow movements are shown as dashed-gray lines and fast as full lines. **(e)** The development of average slopes during low visual uncertainty over time from cursor jump/vision onset. Simulated data is shown in gray, data of experiment 1 in black, and data from experiment 2 in red. Dashed lines correspond to slow and full lines to fast movements.

Figure 7d depicts the temporal evolution of the simulated slopes at 100, 200, 300, and 400ms following the cursor jump (purple lines shown in Fig. 7a). The development of the slopes qualitatively resembles the results shown in Figure 3b and illustrates a similar effect of visual uncertainty and movement speed. Figure 7e depicts a direct comparison of model and data for slopes during the condition with low visual uncertainty. The full and dashed gray line show that it takes ∼300ms following the cursor jump for the slope during the fast simulation to converge to the value of -1, whereas during the slow simulation it takes an additional 100ms to reach the same value. This pattern looks similar when comparing the fast and slow movements in Experiment 1, however there are some important differences between the model simulations and the data. Firstly, while the model starts to exhibit negative slopes at around 100ms following the visual perturbation, we only observed slopes significantly different from 0 at ca. 300ms after vision onset in Experiment 1. This shift in time might be due to the fact that our model does not include time delays in the observed visual feedback. Secondly, the slopes we measured in our experiments converged to a value between -0.5 and -0.75, meaning that the cursor shift was not fully compensated. This divergence between the model and our data might have resulted from the fact that the visual feedback was only briefly presented in the experiments.

In summary, our simulations show that the effect of visual uncertainty can be explained by an optimal state estimator that performs a continuous integration of sensory feedback and internal predictions, while the increase in movement speed induces a change in the control policy that maps state estimates onto motor commands. The results of our model mirror our empirical observations. Specifically, the finding that the contribution of visual feedback increased with viewing duration in Experiment 2 suggests that visual information was indeed accumulated and integrated continuously by the state estimator during the movement. Further, the speed-dependent modulation of forces and EMGs supports that the increased visual feedback response during fast movements was mitigated by an increase in control gains.

## Discussion

The present study aimed to investigate the influence of visual uncertainty on feedback corrections to combined visual and mechanical perturbations. To study the dynamics of the multisensory feedback responses, we varied both the movement time and the presentation duration of the visual feedback during the movement. Our results show that feedback corrections scaled with visual uncertainty and increased during faster movements. Further, we observed that extending the visual feedback duration during slow movements increased its contribution, leading to comparable levels of visual influence during slow and fast movements. We then leveraged a computational model to show in theory that there were two separable components underlying the observed behavior: the first is the integration of visual signals into the motor correction which depends on sensory uncertainty, the second is the modulation of control gains with movement speed. Thus, we conclude that dynamic integration of vision and proprioception in our task was driven by a continuous accumulation of sensory evidence for state-estimation. This estimate in turn interacted with the control policy which scaled with movement time.

Previous work has shown that humans behave close to optimal observers when combining perceptual priors and sensory evidence (Körding and Wolpert, 2004). Specifically, Izawa and Shadmehr (2008) showed that vigor of responses to target jumps depended on the change in relative uncertainty from the first to the second target location. They further demonstrated that the time course of these uncertainty-modulated responses could be predicted by a Kalman-filter based integration of internal priors and delayed sensory feedback, which led to a gradual convergence of the estimated target position towards the new target position. Our findings extend this model to multimodal perturbation responses by showing that feedback responses to force perturbations were modulated by visual uncertainty in a manner predicted by a Kalman filter.

In our second experiment, we observed that an extension of the visual feedback duration resulted in an increased contribution of vision to movement corrections. This observation supports the idea that visual evidence was integrated over the time course of the movement in a process resembling evidence accumulation described during decision-making. Specifically, drift-diffusion models predict that the rate at which the decision variable increases towards a decision threshold depends both on the reliability of the sensory input as well as on the duration during which the sensory evidence is observed (Ratcliff et al., 2016). In the context of reaching control, a stable Kalman filter predicts that estimation errors decay exponentially following a perturbation, and the rate of this decay is determined by the sensory feedback uncertainty. Further, by iteratively collecting information about the system state at each time point, a Kalman filter effectively implements a process similar to evidence accumulation. Hence, changing the duration of the visual feedback likely resulted in more information about the visual stimulus being accumulated by the state estimator, and hence a larger reliance on vision. Future work may look more systematically at the influence of a larger variety of feedback presentation times on state estimation during movement.

More recent studies have introduced the urgency gating model as alternative to drift-diffusion accounts of decision-making. Contrary to the drift-diffusion model, urgency gating does not assume that sensory evidence is accumulated over time and has been able to explain the influence of transient increases in sensory evidence applied at different times during the decision-making process (Carland et al., 2016). Instead, this model proposes that sensory evidence is passed through a low pass filter with a short time constant and then multiplied with an urgency signal that grows over time (Cisek et al., 2009). Interestingly, a growing urgency signal will effectively act like a gain applied to the current sensory evidence, which closely resembles the influence of an increase in control gains due to movement speed as observed in our results. According to Optimal Feedback Control, motor commands are computed by mapping these control gains onto the estimated state of the system. An increase in movement speed is accompanied by a rise in urgency to respond to the perturbation which has been shown to result in an increase in control gains (Crevecoeur et al., 2013; Dimitriou et al., 2013; Poscente et al., 2021). In our experiments this resulted in a transient phase during which the visual compensation seemed larger during fast movements, while the actual estimation error was not influenced by movement speed but was simply mapped onto a different control function. Importantly, urgency gating models of decision-making assume that only novel evidence should influence the decision process which is implemented by assuming a leaky integration process. In our study the hand was moving during the presentation window of the visual feedback, hence each moment in time presented novel evidence about the hand location which was integrated by the state estimator. While resolving the debate between evidence-accumulation and urgency-gating accounts of decision-making is beyond the scope of our study, we demonstrate here that multisensory feedback corrections are influenced by both of these processes. In particular, we suggest that evidence accumulation is performed by the state estimator whereas urgency signals modulate the control policy with which the sensorimotor system responds to incoming feedback signals. Thus, these two processes can be well separated within the context of reaching control.

Our results provide additional support that decision-related evidence accumulation and movement execution cooccur in time, and that decision variables are continually transferred to motor areas during movement as has been suggested by previous work. For example, Selen and colleagues (2012) demonstrated that the accumulated sensory evidence during a decision process was continuously transferred to the motor system during movement preparation by showing that long-latency reflex gains scaled with the accumulated evidence. Further, the occurrence of changes of mind after the onset of a movement demonstrates that the decision process overlaps with movement execution (Visser et al., 2023). Other studies have shown that speed constraints imposed on decision making can influence the speed of subsequent movements and vice versa (Carsten et al., 2023). In line with such bi-directional influences between decision-making and movement control, Kelly and O’Connell (2013) showed that a centroparietal positivity component in an EEG study scaled both with coherence of a random dot motion stimulus as well as with the reaction time of the subsequent decision. The authors linked this effect to a reduction in alpha-band power preceding faster responses which is commonly interpreted as an increase in attention to the stimulus. Although speculative, a similar process might underly the increase in response to visual feedback during faster movements we observed in our data.

The existence of varying processing delays within different sensory modalities has inspired the proposal that initial movement corrections might rely on intramodal estimates until multimodal state estimates become available later on during processing (Oostwoud Wijdenes and Medendorp, 2017). A recent study has shown that initial responses to visual and proprioceptive perturbations exhibited additive effects whereas interactions between these feedback modalities only became visible around 100ms after the onset of responses to visual feedback (Keyser et al., 2023). While the authors interpreted this as evidence against a Kalman filter based integration, we demonstrated here that a linear state-estimator indeed predicts such additive contributions of proprioceptive and visual errors to feedback responses. Nonetheless, a Kalman filter is a description of the behavioral output and does not make assumptions of how the brain produces this behavior. Previous work has shown that different pathways exist both within the proprioceptive and visual modalities (Cross et al., 2019; Pruszynski et al., 2010; Scott, 2016), such that the resulting neural command sent to the muscles is a combination of both independent and combined pathways. The question is where in the brain information from these different pathways is combined for multimodal processing. For example, Bakola and colleagues (2010) used fluorescent tracers in cynomolgus monkeys to show that neuronal populations in parietal cortex receive both visual and proprioceptive inputs specifically linked to limb positioning in space. Further, a recent study has shown that limb afferent feedback and visual information about limb and target location converge on similar neuron populations in primary motor cortex of monkeys (Cross et al., 2021, preprint). Given the latencies that we observed for the visual feedback in our study (ca. 180-220ms), these responses necessarily relied both on separate and combined neural pathways. Hence, an important question for neurophysiological studies is to investigate how multimodal pathways complement separate sensory processing to produce a behavioral output that matches the prediction of a Kalman filter.

In summary, our results show that multimodal feedback responses during movement not only depend on sensory uncertainty, but are further influenced by movement speed and visual feedback duration. Importantly these two influences can be respectively linked to the urgency as a feature of the control policy, and to evidence accumulation over time as a property of the state estimator. From this perspective, these two components of behavior observed across decision-making and motor control tasks can be dissociated and attributed to well-defined computational operations of the sensorimotor system.

## Acknowledgements

AH was supported by an FRS-FNRS FRIA PhD grant (number: FC 036239). FC is supported by an FRS-FNRS grant (number: 1.C.033.18). This work was additionally supported by a Concerted Research Action of UCLouvain (ARC, “coAction”).

